# RapCluster: Bridging the Reproducibility Gap in Clustering Analysis

**DOI:** 10.64898/2026.04.14.718399

**Authors:** Ahmad Lutfi, Robert Warneke, Lutz Fischer, Juri Rappsilber

**Author notes:** Corresponding author: Juri Rappsilber.

## Abstract

Clustering is ubiquitous across science, yet a text-mining audit of 736,399 open-access articles identified as using clustering (2000-2025) reveals common practice leaves key parameters undocumented or untuned, contributing to the reproducibility crisis in science. We developed an interactive web platform featuring 11 widely adopted clustering algorithms to enable transparent clustering analysis and reporting, aligning practical use with best practices in computational research.

**Code availability:** The browser-based clustering analysis platform RapCluster is available for download at https://github.com/lutfia95/RapCluster under MIT License and accessible at https://rappsilberlab.org/rapcluster/ The web version is capacity-limited to 8 GB of memory (ca. 12,000 candidates with 130 features). All source codes used for the analysis are included with the publication.

## Main Text

Researchers in fields from genomics to social sciences rely on clustering to discover patterns in unlabelled data. Unfortunately, this widespread use is accompanied by inconsistent reporting of methodology, which can impede reproducibility. For instance, the very definition of what constitutes a *cluster* is debated^1^, and clustering often yields arbitrary groupings that depend on initial conditions or features. Prior comparative studies in biomedicine have highlighted the need for careful algorithm selection and validation^2^. Similar concerns have been raised in the evaluation of single-cell RNA-seq clustering^3^ and in the application of clustering to social network data^3,4^. However, the community lacks a broad understanding of how clustering methods are actually used in practice in recent literature.

We systematically reviewed open-access research publications from 2000 to 2025 that reported the use of clustering algorithms, using a text-mining pipeline applied to PubMed Central (PMC) full-text JATS XML articles (yielding 736,399 full-text JATS XML articles). Each article was scanned for mentions of clustering methods and annotated using regex-based reporting indicators for *parameters, justification, evaluation*, and *tuning*. In addition, we defined a composite *reporting signals* label to indicate articles in which at least one of these reporting elements was not explicitly detected. These indicators were designed to capture common reporting omissions, including failure to state key algorithm parameters, lack of rationale for algorithm choice, absence of reported cluster-validation procedures, and missing evidence of hyperparameter tuning.

Across the 2000-2025 period, the number of analyzed PMC articles increased markedly, and the proportion of articles with at least one clustering-algorithm match remained high throughout (Figure 1A). “Algorithm match” refers to any article with a clustering-related algorithm signal in the text, including both specific named methods and generic clustering terminology. In general, the clustering algorithm is clearly indicated (90.7% in 2000, 93.1% in 2025). However, explicit textual reporting of key methodological elements remains incomplete in many articles, increasing only modestly from 0.0% in 2000 to 3.2% in 2025 (Figure 1B). At least one key reporting element was not explicitly detected in most articles. Among the individual categories, missing parameters was the most frequent issue (80.2%), followed by missing tuning (78.3%) and missing evaluation (71.8%), whereas missing justification was less common (22.5%). These results indicate that important methodological details were often not explicitly reported in article text (Figure 1B).

**Figure 1.**
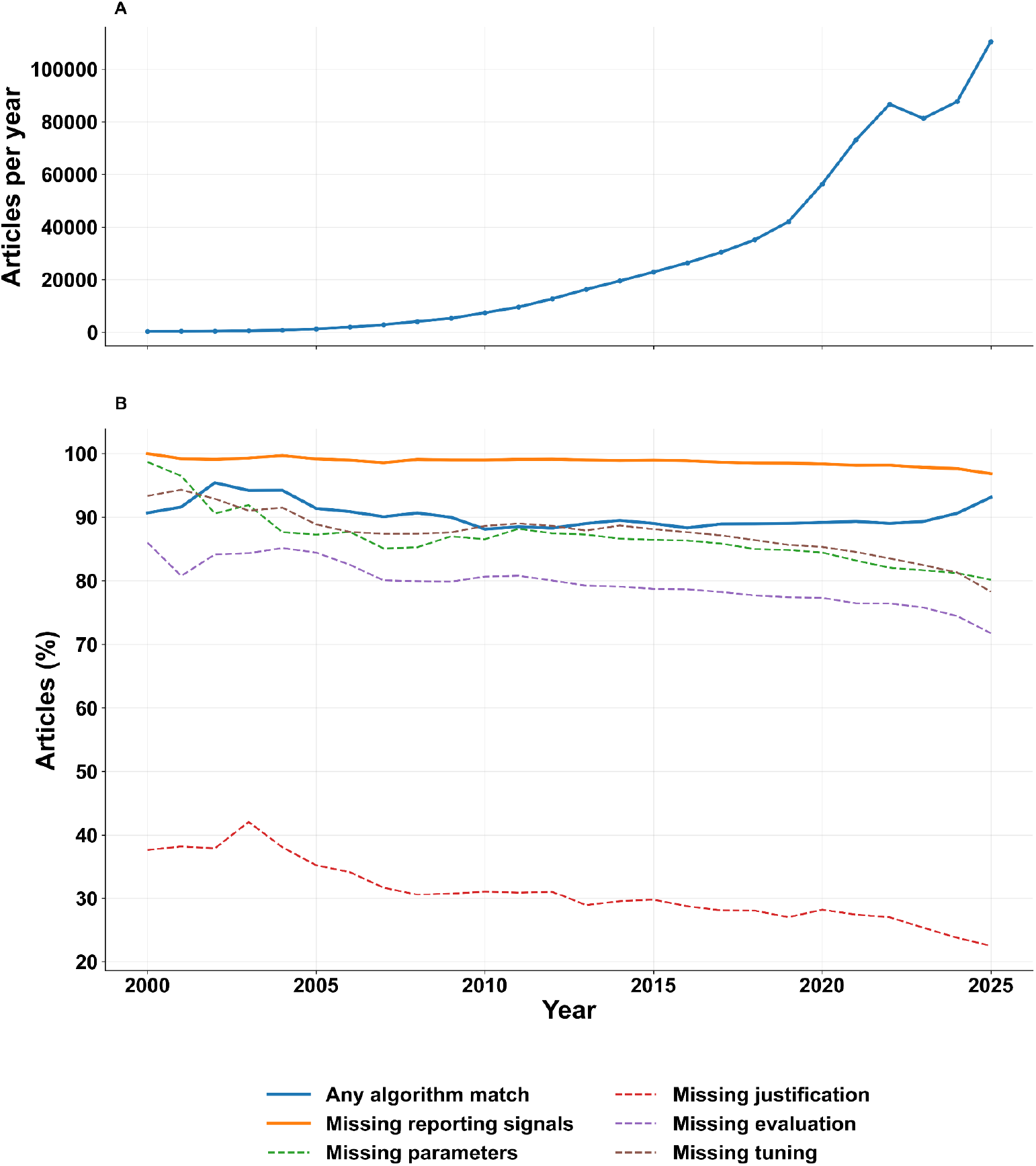
Longitudinal trends in clustering-method reporting across PMC articles, 2000-2025. **A**, Total number of analyzed PubMed Central open-access full-text articles per year. **B**, Year-wise proportions of articles with any clustering-algorithm match (including generic clustering terms) and with missing reporting signals, including missing parameters, missing justification, missing evaluation, and missing tuning. Across the full 2000–2025 window, corpus size increased substantially, while missing reporting signals remained common despite high rates of algorithm detection in recent years. Percentages were computed at year level from the text-mining summary.

This gap is concerning, as evaluating clustering results is essential to ensure that identified patterns are meaningful and not artefacts^1^. In particular, the majority of studies did not describe any hyperparameter tuning or sensitivity analysis for their clustering (for example, simply running *k*-means with a single choice of *k* without justification). Neglecting to explore parameter settings can lead to suboptimal or biased results, since clustering outcomes can vary significantly with different parameter choices depending on data type^5^. Taken together, the persistently high rates of omissions across all categories, including missing evaluation and missing tuning, suggest that explicit textual reporting of key clustering-related methodological details remains incomplete in many articles. An interactive supplementary HTML summary provides the complete overview of all metrics generated in the 2000-2025 text-mining analysis, including corpus-level statistics, annual trends, and category-specific reporting signals.

We additionally performed an algorithm-stratified analysis to assess whether reporting patterns differed across clustering methods. Reporting patterns were not uniform across clustering methods; instead, the frequency of missing reporting signals varied substantially by algorithm, both in pooled analyses and across publication years (Supplementary Fig. S2). For example, parameters were missing in 93.0% of cases when OPTICS was used for clustering (n = 75,105). In contrast, parameters were missing for k-means in 47.7% of cases (n = 51,759). This discrepancy, particularly given the less prominent parameter visibility within OPTICS implementations compared to k-means, indicates that reporting compliance is strongly influenced by the practical ease of documenting parameters.

Recognizing the systematic differences in reporting practices and the urgent need for tools that standardize and simplify transparent methodological documentation, we developed an interactive browser-based tool, RapCluster, to help avoid in future the shortcomings observed in the literature.

Users can upload their own datasets and apply a variety of clustering (e.g., KMeans, DBSCAN, HDBSCAN, Gaussian Mixture Models) and dimensionality reduction algorithms (PCA, t-SNE, UMAP). Default parameters are provided for each method, but users can flexibly tune them to explore the impact of algorithmic choices on their data. Crucially, the platform guides the user through important steps that are often overlooked. For example, when a user selects an algorithm, the tool prompts for key parameters (e.g., *k* for k-means) and provides short explanations indicating which ones are recommended for fine-tuning. In this way, users are encouraged to consider multiple options rather than defaulting to a single run.

Another core feature is the built-in evaluation of results. The application will compute and display relevant clustering validation metrics, including silhouette score. Visualizations are generated dynamically to help interpret the clusters (e.g., cluster network). Importantly, the tool doesn’t stop at analysis: it automatically generates text that the researcher can use in a publication. This text is context-aware, it includes the chosen algorithm name, key parameter values, and the evaluation metrics, phrased in complete sentences. By providing this auto-generated text, the tool aims to lower the barrier to thorough reporting, even researchers unfamiliar with clustering will have a template for how to describe their analysis properly. We anticipate that this feature will especially help improve the “Methods” sections of papers and ensure key details are not omitted.

We envision this interactive platform as both an educational and practical resource. To illustrate its usage, Figure 2 showcases the tool’s output on an example dataset from^6^ detailing the growth of *Bacillus subtilis* genome-scale deletion mutants under various growth media conditions, which provided insights into gene functions. In our tool, we uploaded this dataset (mutant strain) and applied clustering to identify groups of mutants with similar fitness profiles. These data are provided for demonstration purposes only, without analytical intent. The tool’s output included a caption describing these clusters and noting the evaluation metrics, as well as a methods paragraph detailing the clustering procedure. We designed the interface to be accessible to non-experts, with pop-up explanations and tips (including short video walkthroughs). Our hope is that by making best practices the path of least resistance, the tool will foster more consistent and transparent use of clustering.

**Figure 2.**
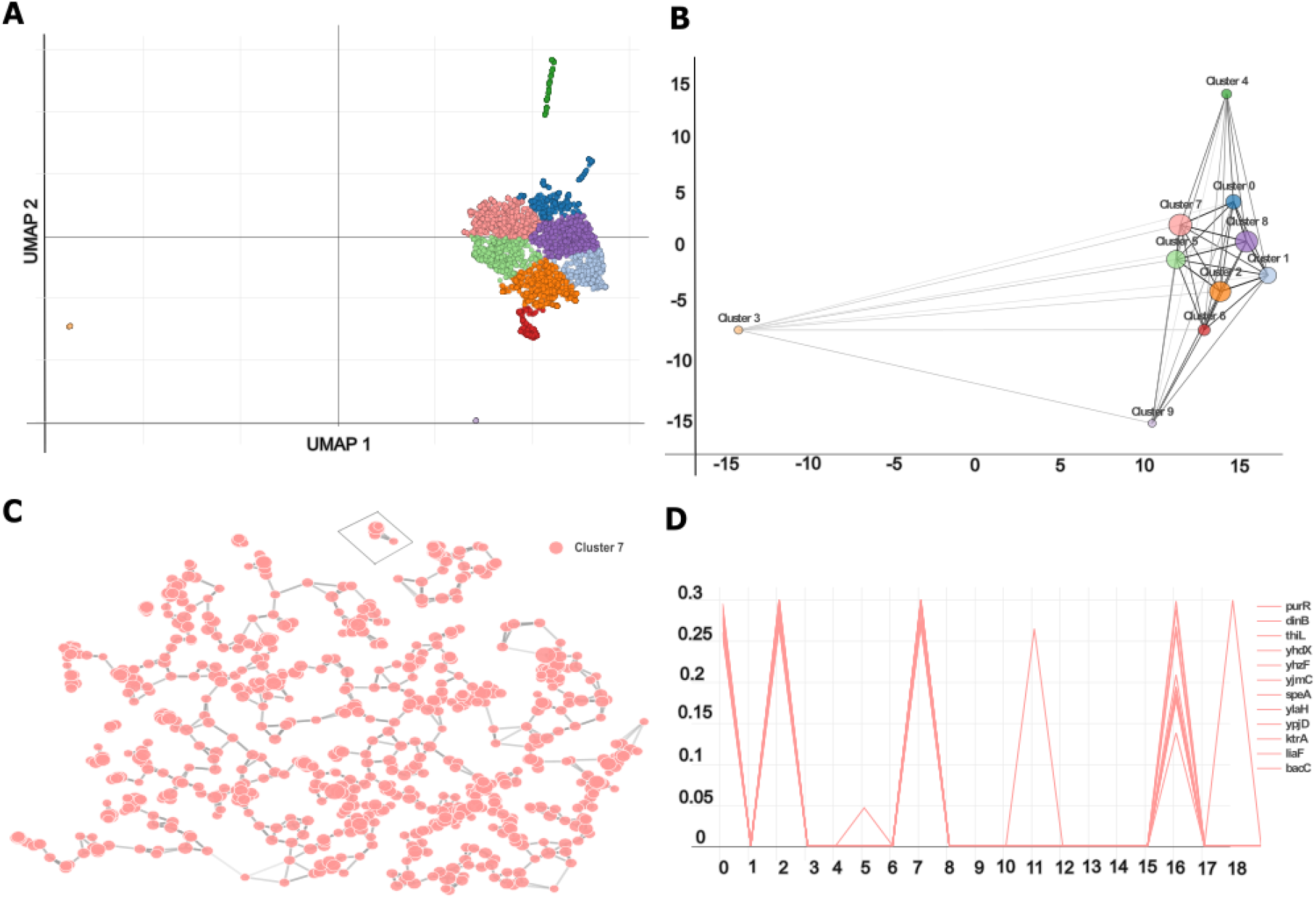
Demonstration of clustering analysis using our interactive web application. **A**, Two-dimensional visualization of Bacillus subtilis genome-scale deletion mutant data ^6^, clustered into distinct groups using unsupervised learning. **B**, Clusters connection network weighted with Euclidean distances. **C**, High-dimensional view of the selected cluster (7). **D**, Visualization of the selected sub-cluster is provided through line plots to identify shared patterns between intensity-profiles indexed from 0 to 19.

In conclusion, while clustering algorithms are more popular than ever, crucial aspects of methodology are frequently under-reported. This underscores ongoing challenges in the field, reminiscent of broader concerns in machine learning about reproducibility and transparency^7^, and highlighting the importance of active measures to support the FAIR data principles (Findable, Accessible, Interoperable, Reusable). By openly quantifying these issues, we aim to raise awareness. More immediately, RapCluster offers researchers a way to actively close the gap between actual and best practice. It serves as a platform for trying out clustering methods and as a safety net that catches omissions in reporting. All components of RapCluster are open-source, so the community can contribute new algorithms or features. As the use of clustering continues to grow (Figure 1), we aim to ensure the platform remains a valuable, evolving resource for the scientific community.

## Supporting information

Supplemental analysis (html)

## Acknowledgements

This work was supported by the Deutsche Forschungsgemeinschaft (DFG, German Research Foundation) under Germany’s Excellence Strategy - EXC 2008 - 390540038 - UniSysCat and the European Research Council (ERC) under the European Union’s Horizon Europe research and innovation programme (grant agreement No. 101119142 - TransFORM).

## Methods

### Systematic audit of clustering‐parameter reporting Publications acquisition

We programmatically queried PubMed Central (PMC) for open-access articles published between 2000 and 2025 using the NCBI E-utilities^8^ interface through Biopython’s Bio.Entrez^9^ module. Searches were performed against the pmc database using a Boolean query targeting clustering-related terminology, including both generic and algorithm-specific terms, together with the PMC open-access filter and a year-specific publication-date restriction. Publication years were iterated across the full 2000–2025 study window, and matching PMCIDs were retrieved as JATS XML records using *efetch* and stored in year-specific directories for downstream processing. We set Entrez.email and Entrez.api_key in accordance with NCBI guidance, respected rate limits, and implemented retry logic with backoff and HTTP error handling. To improve throughput for I/O-bound retrieval, downloads were parallelized using Python’s concurrent.futures framework, with progress tracking via *tqdm*. The acquisition workflow relied on NCBI E-utilities, Biopython/Entrez, the PMC Open Access subset, and JATS XML.

### Text Mining and Relevance Determination

Following XML retrieval and parsing, we applied an automated text-mining pipeline to identify clustering-method mentions and reporting signals in the article body text. The implementation was written in Python 3.11 and relied primarily on *pandas* for structured data handling and Python’s built-in re module for regular-expression matching. The pipeline operated on normalized body text, and all article-level signals were defined as regex-based extractions from that text.

Each article was screened for clustering-algorithm mentions and for four reporting categories: missing parameters, missing justification, missing evaluation, and missing tuning. Distinct algorithm matches were recorded in the field *algorithms_found*; articles with at least one detected algorithm mention contributed to the yearly metric for articles with any algorithm match. For reporting indicators, *missing_params* denoted absence of a parameter-related regex match, *missing_justification* absence of a justification-related match, *missing_evaluation* absence of an evaluation-related match, and *missing_tuning* absence of a tuning-related match. We further defined a composite indicator, *missing_reporting_signals*, which was assigned when at least one of the four missing-category indicators was present. Thus, in this study, “missing” refers to absence of an explicit regex-detectable reporting signal, rather than proof of methodological misuse.

Label assignment was performed using curated sets of regular expressions tailored to algorithm detection and to each reporting category. These regex banks targeted both algorithm-specific terminology and conceptual reporting indicators, including terms related to parameter specification, rationale for method choice, cluster evaluation, and tuning or model selection. Articles could receive multiple missing-category indicators simultaneously. To reduce false positives, patterns were designed to capture both exact keywords and contextual phrases. The full operational definitions and embedded regex banks are provided in the Supplementary Materials and in the supplementary HTML summary and a worked example of the text-mining workflow.

### Algorithm-stratified reporting analysis

To assess whether reporting patterns differed across clustering methods, we performed a secondary algorithm-stratified analysis using the article-level outputs of the main text-mining pipeline. For each article, the primary pipeline provided regex-derived fields including *algorithms_found, missing_params, missing_justification, missing_evaluation, missing_tuning*, and the composite *missing_reporting_signals*, where “missing” denotes absence of a reporting signal in normalized body text. Detected algorithm mentions were normalized to a predefined set of standardized clustering labels using case-insensitive regular expressions. This mapping included both classical clustering methods and related graph- and single-cell-oriented clustering terms. Because all detected hits were retained, a single article could contribute to multiple algorithm labels when more than one clustering method was mentioned. For each standardized algorithm label, we quantified the number of associated articles and the proportions showing missing parameters, missing justification, missing evaluation, missing tuning, and missing reporting signals overall. Summaries were generated both by publication year and across the full pooled dataset. To avoid unstable year-specific estimates, algorithm-by-year summaries were retained only for labels represented by at least 50 articles in the corresponding output. For the pooled analysis, algorithm-specific missing-reporting rates were compared against the overall baseline among articles with at least one detected algorithm mentioned. Differences in overall missing-reporting proportions were evaluated using two-proportion *z*-tests, and the resulting *p*-values were adjusted for multiple testing using the Benjamini-Hochberg false-discovery-rate procedure.

### Interactive web application for clustering parameter tuning Architecture

We implemented a Python backend exposing two HTTP endpoints-GET */api/algorithms* and POST */api/cluster* and a React/Plotly^10,11^ frontend that drives parameter selection, submission, and visualization. The frontend uses *react-plotly.js* for interactive charts and calls the backend, passing the uploaded file and the chosen reduction/clustering parameters. An overview of the application is provided in the Supplementary video.

### Data ingestion and schema

Users upload table file (*.TSV), specify a name field (node identifier) and the first column index, where the clustering values start. The backend returns: (i) cluster data per item: name, cluster, 2D coordinates (x,y) from the selected reduction; (ii) metrics: *n_clusters, silhouette_score, calinski_harabasz_score* and *davies_bouldin_score*^*10,12*^; (iii) cluster colors; a color per cluster label for consistent plotting.

### Parameterization interface

The UI lists available clustering algorithms and dimensionality-reduction methods from GET */api/algorithms*, along with per-method parameter definitions (type, default, ranges, allowed options). Users can keep defaults or supply custom values (including nullable None where allowed). All selections are serialized as JSON and sent with the uploaded data to POST */api/cluster*.

### Algorithms implemented

The app exposes, KMeans, MiniBatchKMeans, AgglomerativeClustering, AffinityPropagation, MeanShift, SpectralClustering, DBSCAN, OPTICS, BIRCH, GaussianMixture, and HDBSCAN. Implementations follow scikit-learn (all listed except HDBSCAN) and hdbscan^13^ (for HDBSCAN).

### Dimensionality reduction

The app supports PCA, t-SNE, and UMA. PCA and t-SNE follow scikit-learn; UMAP follows umap-learn^13,14^. Typical parameters exposed include *n_components, n_neighbors, min_dist, perplexity, learning_rate*, and *random_state*.

### Scoring and reporting

For every run, the backend computes Silhouette (higher is better), Calinski–Harabasz (higher), and Davies–Bouldin (lower), returned with the number of clusters and runtime. Metric definitions and APIs follow scikit-learn’s *sklearn.metrics*.

### Network construction (centroid graph)

From the 2D embedding, we compute a centroid per cluster (mean of *x,y*) and build a complete graph over centroids. Edge weights are Euclidean distances^15^ in the 2D reduced space; line width is inversely related to distance and opacity/greyscale encodes relative length. Node size scales with cluster size; node color is the cluster color returned by the backend.

### Subnetwork extraction (intra-cluster k-NN graph)

Selecting a cluster builds an intra-cluster *k-nearest-neighbors* graph (default top 5 neighbors per node, user-adjustable). Distances are Euclidean in the same 2D space. Node size reflects degree; edges are drawn with width ∼1/distance and opacity scaled by relative distance to improve readability. Users can lasso-select nodes to render *log10* intensity profiles for the selected items. All behaviors are implemented in the frontend handlers (handleSubnetworkClick, handlePointsSelected).

### Visualization layer

All plots use Plotly via *react-plotly.js* (scatter for embeddings, centroid networks, and intra-cluster subnetworks; line plots for selected profiles). Figures can be exported to SVG via the toolbar.

### Reproducibility & defaults

Where applicable, *random_state* is exposed (PCA/t-SNE/UMAP and algorithms that support seeding) to ensure reproducible runs. All default values and constraints (type, min/max, step, options, nullable) are served by the backend and echoed back in the UI’s *“Method Details”* panel for provenance.

## Notes

https://rappsilberlab.org/rapcluster/

### Competing Interest Statement

The authors have declared no competing interest.

