## Supplemental PDF for "RapCluster: Bridging the Reproducibility Gap in Clustering Analysis"

### Supplementary Materials for “RapCluster: Bridging the Reproducibility Gap in Clustering Analysis”

**Supplementary Methods**: **Operational Definitions**

To operationalize the methodological assessment described in the main text, we designed a series of regular expression (*regex*) patterns targeting specific linguistic markers of clustering methodology. These patterns enabled systematic and reproducible detection of methodological omissions or misuses across abstracts and method sections. Five main categories of labels were used: *Missing Reporting Signals*, *Missing Parameters*, *Missing Justification*, *Missing Evaluation*, and *Missing Tuning*. Each category was associated with a curated set of *regex* rules that capture either the explicit mention of relevant terms (e.g., *silhouette score*, *grid search*) or the absence thereof. The following table (Supplementary Table S1) provides a detailed mapping of labels to regex patterns and explanations, ensuring transparency and reproducibility of the text mining approach.

| **Label** | **Regex Patterns and Explanation** |
|---|---|
| **Missing Parameters:** was assigned when no match was detected in the *params_found* regex bank. This bank was designed to capture explicit mentions of clustering-related parameter settings in normalized article body text. | Patterns targeted commonly reported clustering parameters and closely related configuration terms, including:<br><br>- Number of clusters / cluster count (*n_clusters*, “number of clusters”, *k* = ...) - captures explicit specification of cluster number, particularly for partition-based methods such as *k*-means.<br>- Initialization settings (*init* = ..., *n_init* = ...) - captures explicit initialization choices and restart settings, especially relevant to k-means-type algorithms.<br>- Iteration and convergence parameters (*max_iter*, *tol*) - captures explicit |

| | |
|---|---|
| | optimization or stopping criteria.<br>- Density-based clustering parameters (*eps*, *min_samples*, *min_cluster_size*, *xi*, *cluster_method*) - captures key settings used in DBSCAN-, HDBSCAN-, and OPTICS-related workflows.<br>- Hierarchical-clustering parameters (*linkage* = ward\|complete\|average\|single) - captures explicit agglomeration settings.<br>- Additional explicit parameter notations such as parameter assignments and method-specific configuration keywords - captures other structured parameter mentions included in the regex bank. |
| **Missing Justification:** *missing_justification* was assigned when no match was detected in the *justification_found* regex bank. This bank was designed to capture explicit textual rationale for choosing clustering settings such as cluster number, resolution, or related decision rules in normalized article body text. | Patterns targeted explicit justification language and common rule-based selection criteria, including:<br><br>- Rule-based cluster-selection terms such as *elbow*, *knee*, *gap* statistic, and silhouette analysis/score - captures explicit statements that cluster number or resolution was selected using a recognized decision criterion.<br>- Direct choice statements such as chosen/selected/determined/set based on, we chose/selected/set/determined, and was chosen/selected/determined/set - captures explicit author statements describing how a clustering setting was selected.<br>- Explicit determination of cluster number such as to determine the number of clusters and optimal k / number of clusters / resolution - captures text indicating that a parameter choice was actively derived rather than merely reported.<br>- Empirical or literature-consistency language such as empirical/empirically and consistent with previous/prior - captures justification based on empirical |

| | |
|---|---|
| | - assessment or alignment with prior work. |
| **Missing Evaluation:** *missing_evaluation* was assigned when no match was detected in the *evaluation_found* regex bank. This bank was designed to capture explicit mentions of cluster-validation metrics or model-diagnostic criteria in normalized article body text. If *evaluation_found = 0*, then *missing_evaluation = 1*. | Patterns targeted commonly reported internal, external, stability-based, and model-selection evaluation terms, including:<br><br>- Internal validation metrics such as silhouette, Davies-Bouldin, and Calinski-Harabasz - captures explicit mention of standard cluster-quality metrics.<br>- Dispersion and variance criteria such as within-cluster sum of squares (WCSS) and between-cluster variance- captures explicit optimization or separation criteria used in cluster assessment.<br>- Stability-based evaluation terms such as cluster stability, stability analysis, and bootstrap … cluster - captures explicit mention of robustness or resampling-based validation.<br>- External validation metrics such as adjusted Rand index (ARI), normalized mutual information (NMI), adjusted mutual information (AMI), Fowlkes-Mallows, and Jaccard - captures comparison against reference labels or alternative partitions.<br>- Model-diagnostic criteria such as BIC, AIC, log-likelihood, and modularity - captures model-selection or graph/community-evaluation terminology included in the evaluation regex bank. |
| **Missing Tuning:** *missing_tuning* was assigned when no match was detected in the *tuning_found* regex bank. This bank was designed to capture explicit hyperparameter-search and model-selection language in normalized article body text. If *tuning_found = 0,* then *missing_tuning = 1*. | Patterns targeted explicit tuning and search terminology, including:<br><br>- General hyperparameter-tuning language such as hyperparameter tuning, hyperparameter optimization/optimisation, and parameter sweep - captures explicit statements that parameter search was performed.<br>- Systematic search strategies such as grid search and random/randomized |

| | |
|---|---|
| | search - captures structured search over parameter space.<br>- Optimization-based tuning approaches such as Bayesian optimization/optimisation - captures model-based hyperparameter search.<br>- Optimization libraries such as Optuna, Hyperopt, and skopt - captures named software frameworks for hyperparameter optimization.<br>- Validation-based tuning language such as cross-validation, CV, and nested cross-validation - captures explicit tuning or selection procedures based on repeated validation.<br>- Explicit tuning/action phrases such as tuned/optimized/optimised/selected using and we tuned / we optimized / we optimised - captures direct author statements that parameter tuning was performed. |
| **Missing Reporting Signals:** *missing_reporting_signals* was defined as a composite indicator summarizing the four reporting categories. It was assigned when at least one of the following article-level indicators was positive: *missing_params*, *missing_justification*, *missing_evaluation*, or *missing_tuning*. In logical form: *missing_reporting_signals = missing_params OR missing_justification OR missing_evaluation OR missing_tuning* | Composite indicator assigned when at least one of the following was positive: *missing_params*, *missing_justification*, *missing_evaluation*, or *missing_tuning*. In this study, it denotes absence of at least one regex-detectable reporting signal, rather than confirmed methodological misuse. |

**Supplementary Table S1.** Detailed text-mining criteria and pattern explanations for reporting-signal assignment. Regex-based criteria were applied for the reporting categories Missing Parameters, Missing Justification, Missing Evaluation, and Missing Tuning. The composite variable Missing Reporting Signals was assigned when at least one of these four category-specific indicators was positive. In this study, these labels denote absence of regex-detectable reporting signals in normalized article body text, rather than confirmed methodological misuse. Full implementation details are provided in the accompanying code and supplementary HTML summary.

**Supplementary Methods**: **Example Text-Mining**

**Paper text** [1]**:**
*…… We used the hierarchical clustering (with Ward linkage = 3) as the baseline algorithm for this experiment and evaluated the outlier cutoffs from 0.2 to 0.5, with a step size of 0.1……*

*…… Instead, the method returns as the optimal number of clusters the value at which the highest Silhouette score was obtained; in this case, the suggested optimal number is 2 cluster……*

*…… Even when all metrics reach their best value with different numbers of clusters, the difference in the results with respect to the best value is small. In fact, the difference of the best silhouette coefficient with respect to the other values is between 3–6%……*

*…… The evaluation of this experiment was done in terms of the three clustering quality metrics. Once these hyperparameters have been set, the second step of hyperparameter tuning uses these values and focuses on finding the optimal number of clusters by using the elbow method……*

*…… However, we require a compromise between the number of clusters and clustering quality as more clusters imply more cross-validations and thus more expensive computation……*

**Workflow output:**

Algorithms_found: hierarchical clustering

Params_found: Ward linkag

Justification_matches: Silhouette score, optimal number of clusters

Evaluation_matches: silhouette coefficient

Tuning_matches: hyperparameter tuning, cross-validation

#### Supplementary Figures

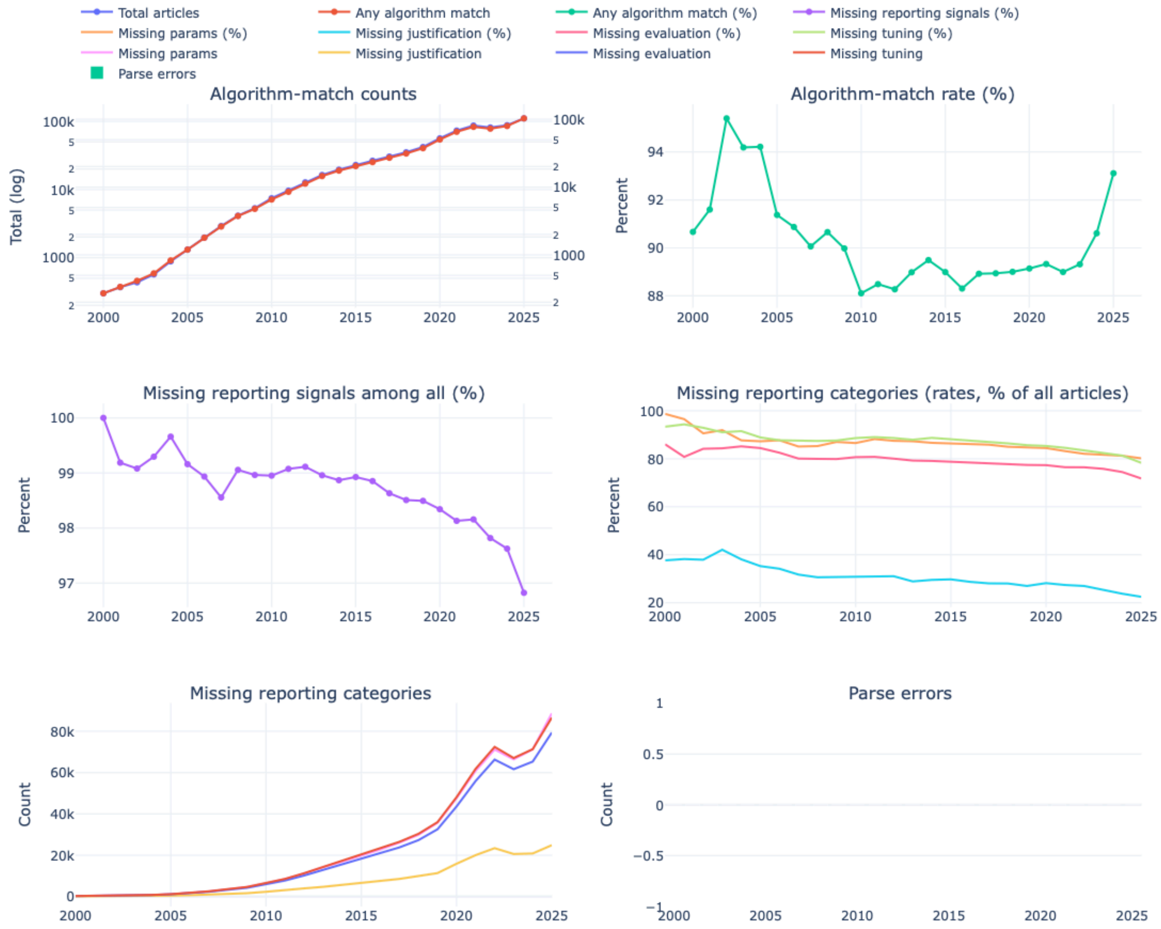


**Supplementary Figure S1. Year-level overview of corpus size, algorithm detection, and reporting signals from 2000 to 2025.** The supplementary HTML summary reports the complete longitudinal output of the text-mining pipeline across the full study period. Shown are total article counts, counts and rates of clustering-algorithm matches, the proportion of articles with missing reporting signals, category-specific rates and counts for missing parameters, justification, evaluation, and tuning, as well as parse-error counts. Together, these panels provide the complete summary of the metrics used in the main text and supplementary analyses. Parse errors define the number of articles, where our parsing system couldn't read them.

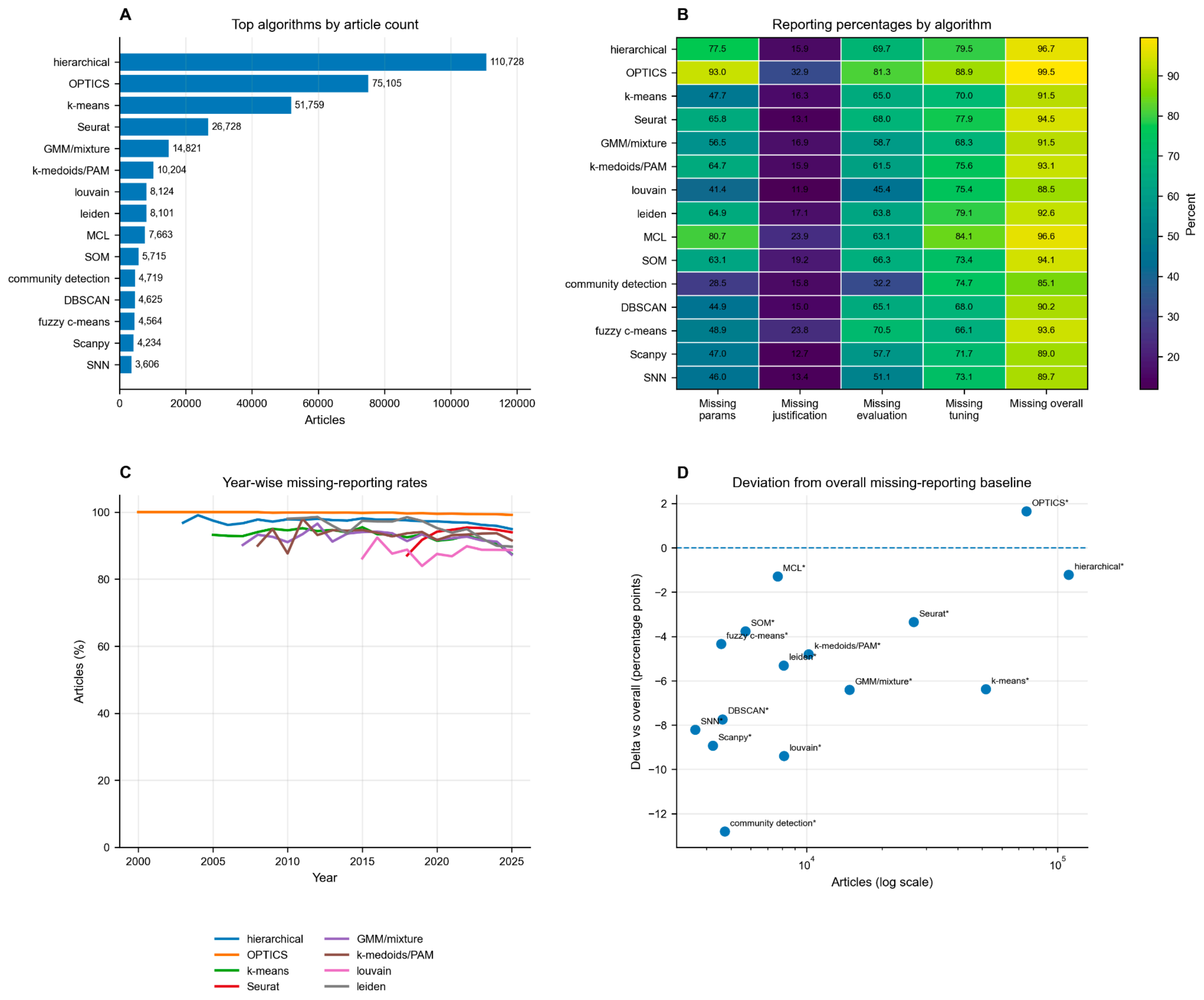


**Supplementary Figure S2. Algorithm-stratified reporting patterns across the corpus. A,** The clustering algorithms most frequently (Top 15) detected in the corpus, ranked by number of associated articles. The plot is not showing the generic cluster match, which we assign as general mentioning of the clustering method. **B,** Per-algorithm percentages of articles with missing parameters, missing justification, missing evaluation, missing tuning, and overall missing reporting signals. **C,** Year-wise trends in overall missing reporting signals for the most frequently represented algorithms. **D,** Deviation of each algorithm's overall missing-reporting rate from the corpus-wide baseline, expressed in percentage points. For each algorithm, the overall proportion of articles with missing_reporting_signals was compared with the corresponding overall proportion among all articles with at least one detected algorithm mention; p-values were obtained using a two-proportion z-test and adjusted by the Benjamini-Hochberg false-discovery-rate procedure. below 0: papers using that algorithm had fewer missing reporting signals than average, whereas above 0 means; papers using that algorithm had more missing reporting signals than average.

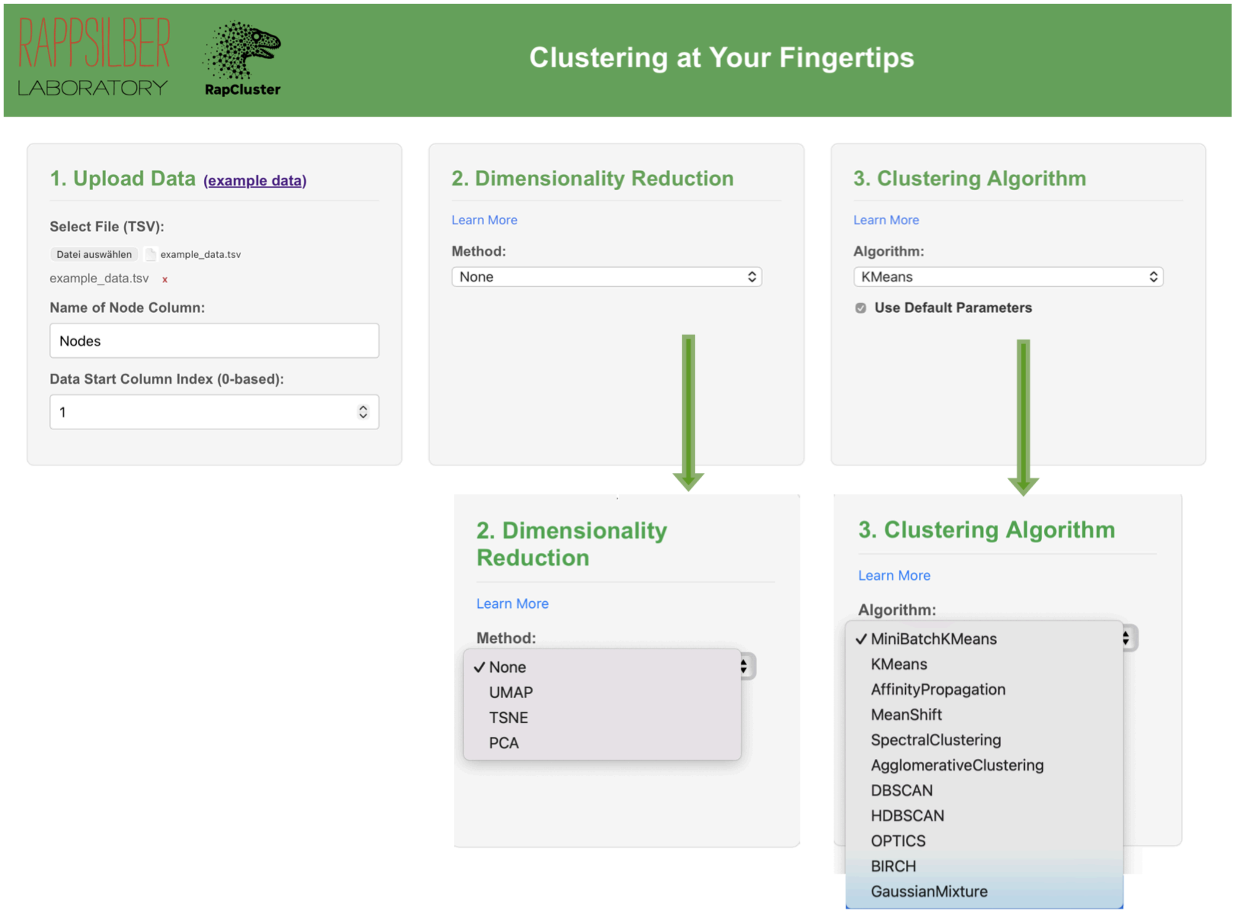


**Supplementary Figure 3.** Overview of the RapCluster web application. The interface allows users to upload their own datasets and apply a wide range of clustering and dimensionality reduction algorithms through an intuitive graphical interface. The tool supports MiniBatchKMeans, KMeans, Affinity Propagation, Mean Shift, Spectral Clustering, Agglomerative Clustering, DBSCAN, HDBSCAN, OPTICS, BIRCH, and Gaussian Mixture models, all with adjustable default parameters. This setup enables researchers to explore clustering outcomes systematically and with greater methodological rigor.
