## Supplemental analysis (html) for "RapCluster: Bridging the Reproducibility Gap in Clustering Analysis": Mining.html

Summary of Datamining (2000-2025)

Year-level summary

Coverage window

2000–2025

Total articles (2025)

110,541

Any algorithm match (2025)

93.11%

Missing reporting signals (2025)

96.82%

Parse errors (2025)

0

Trends

Use the range slider on the bottom panels to focus on specific eras; hover to read exact values.

Composition of missing reporting categories (stacked share, % of all missing flags)

Stacked shares normalize the four missing categories by their per-year sum to show dominance patterns over time.

Methods and definitions

Operational definitions used by the text-mining pipeline, with the exact regex banks embedded.

Snapshot (2025) + metric definitions

| Metric | Value | Definition |
| --- | --- | --- |
| total\_articles | 110,541 | Count of processed XML files assigned to that year. |
| articles\_with\_any\_algorithm\_match | 102,928 | Articles with algorithms\_found non-empty. |
| pct\_with\_any\_algorithm\_match | 93.11% | 100 × articles\_with\_any\_algorithm\_match / total\_articles. |
| articles\_with\_missing\_reporting\_signals | 107,031 | Articles where missing\_reporting\_signals=1. |
| pct\_missing\_reporting\_signals\_among\_all | 96.82% | 100 × articles\_with\_missing\_reporting\_signals / total\_articles. |
| missing\_params | 88,603 | Articles where missing\_params=1 (params\_found empty). |
| missing\_justification | 24,851 | Articles where missing\_justification=1 (justification\_found=0). |
| missing\_evaluation | 79,316 | Articles where missing\_evaluation=1 (evaluation\_found=0). |
| missing\_tuning | 86,536 | Articles where missing\_tuning=1 (tuning\_found=0). |
| parse\_errors | 0 | Rows with non-empty error field; still counted in totals. |

All signals are regex-based extractions from normalized body text; “missing” means “no regex match in that category”.

Field meanings (article-level) — click to expand

Each field includes the exact operational definition and the regex bank used (verbatim).

algorithms\_founddistinct hits from DEFAULT\_PATTERNS["algorithm"]

The pipeline searches the article body text against every regex in DEFAULT\_PATTERNS["algorithm"] with case-insensitive matching.
Each regex contributes all non-overlapping matches it finds. Matched substrings are de-duplicated per article (case-folded) and emitted as a semicolon-joined list.
If the list is non-empty, the article contributes to articles\_with\_any\_algorithm\_match in the yearly summary.

| # | Regex (verbatim) |
| --- | --- |
| 1 | \bk[\s-]?means\b |
| 2 | \bmini[\s-]?batch[\s-]?k[\s-]?means\b |
| 3 | \bk[\s-]?medoids\b\|\bpam\b |
| 4 | \bfuzzy\s\*c[\s-]?means\b\|\bc[\s-]?means\b |
| 5 | \bhierarchical\s+clustering\b\|\bagglomerative\s+clustering\b\|\bdivisive\s+clustering\b |
| 6 | \bward(?:'s)?\b\s+method\b |
| 7 | \bdbscan\b |
| 8 | \bhdbscan\b |
| 9 | \boptics\b |
| 10 | \bmean\s\*shift\b |
| 11 | \bbirch\b |
| 12 | \bspectral\s+clustering\b |
| 13 | \baffinity\s+propagation\b |
| 14 | \bgaussian\s+mixture(?:\s+model)?\b\|\bgmm\b\|\bmixture\s+model\b |
| 15 | \bdirichlet\s+process\s+mixture\b\|\bdpm\b |
| 16 | \bself[-\s]?organizing\s+map\b\|\bsom\b |
| 17 | \bneural\s+gas\b |
| 18 | \bmarkov\s+clustering\b\|\bmcl\b |
| 19 | \bgraph[-\s]?based\s+clustering\b |
| 20 | \bcommunity\s+detection\b |
| 21 | \blouvain\b |
| 22 | \bleiden\b |
| 23 | \bshared\s+nearest\s+neighbor\b\|\bsnn\b |
| 24 | \bphenograph\b |
| 25 | \bseurat\b |
| 26 | \bscanpy\b |
| 27 | \bsc3\b |
| 28 | \bcluster(?:ing\|ed\|s)?\b |


params\_founddistinct hits from DEFAULT\_PATTERNS["params"]

Same matching logic as algorithms\_found, but using DEFAULT\_PATTERNS["params"].
This bank targets explicit parameter mentions (k, n\_clusters, eps, min\_samples, resolution, linkage, etc.).
If params\_found is empty, then missing\_params=1.

| # | Regex (verbatim) |
| --- | --- |
| 1 | \bn[\_\s-]?clusters\b |
| 2 | \bnumber\s+of\s+clusters\b |
| 3 | \b(?:k\|K)\s\*=\s\*\d+\b |
| 4 | \binit\b\s\*=\s\*(?:k[-\s]?means\+\+\|random)\b |
| 5 | \bn[\_\s-]?init\b\s\*=\s\*\d+\b |
| 6 | \bmax[\_\s-]?iter(?:ations)?\b\s\*=\s\*\d+\b |
| 7 | \btol(?:erance)?\b\s\*=\s\*[\deE\.\-]+\b |
| 8 | \beps\b\s\*=\s\*[\deE\.\-]+\b |
| 9 | \bmin[\_\s-]?samples\b\s\*=\s\*\d+\b |
| 10 | \bmin[\_\s-]?cluster[\_\s-]?size\b\s\*=\s\*\d+\b |
| 11 | \bxi\b\s\*=\s\*[\deE\.\-]+\b |
| 12 | \bcluster[\_\s-]?method\b\s\*=\s\*\w+\b |
| 13 | \blinkage\b\s\*=\s\*(?:ward\|complete\|average\|single)\b |
| 14 | \b(?:distance\|affinity)\s+metric\b |
| 15 | \bmetric\b\s\*=\s\*(?:euclidean\|cosine\|manhattan\|cityblock\|chebyshev\|minkowski\|correlation)\b |
| 16 | \bward\b\s+linkage\b |
| 17 | \bcut(?:ting)?\s+the\s+dendrogram\b\|\bcut\s+height\b |
| 18 | \bn[\_\s-]?components\b\s\*=\s\*\d+\b |
| 19 | \bcovariance[\_\s-]?type\b\s\*=\s\*(?:full\|tied\|diag\|spherical)\b |
| 20 | \bregulari[sz]ation\b\s\*=\s\*[\deE\.\-]+\b |
| 21 | \bn[\_\s-]?neighbors\b\s\*=\s\*\d+\b |
| 22 | \bgamma\b\s\*=\s\*[\deE\.\-]+\b |
| 23 | \bdamping\b\s\*=\s\*[\deE\.\-]+\b |
| 24 | \bpreference\b\s\*=\s\*[\deE\.\-]+\b |
| 25 | \bbandwidth\b\s\*=\s\*[\deE\.\-]+\b |
| 26 | \bresolution\b\s\*=\s\*[\deE\.\-]+\b |
| 27 | \bmodularity\b |
| 28 | \bkNN\b\|\bk\s\*nearest\s+neighbors\b |
| 29 | \bneighbors\b\s\*=\s\*\d+\b |
| 30 | \bUMAP\b.\*\b(n[\_\s-]?neighbors\|min[\_\s-]?dist)\b |
| 31 | \bt-?SNE\b.\*\b(perplexity\|learning\s\*rate)\b |


justification\_foundbinary: any hit in DEFAULT\_PATTERNS["justification"]

Set to 1 if at least one regex in DEFAULT\_PATTERNS["justification"] matches the body text; otherwise 0.
This captures explicit rationale for choosing cluster count/resolution/thresholds (elbow/knee, gap statistic, “we chose… based on…”, etc.).
If the value is 0, then missing\_justification=1.

| # | Regex (verbatim) |
| --- | --- |
| 1 | \belbow\b\|\bknee\b |
| 2 | \bgap\s+statistic\b |
| 3 | \bsilhouette(?:\s+analysis\|\s+score)?\b |
| 4 | \b(?:chosen\|selected\|determined\|set)\s+(?:based\s+on\|according\s+to)\b |
| 5 | \bwe\s+(?:chose\|selected\|set\|determined)\b |
| 6 | \bwas\s+(?:chosen\|selected\|determined\|set)\b |
| 7 | \bto\s+determine\s+(?:the\s+)?number\s+of\s+clusters\b |
| 8 | \boptimal\s+(?:k\|number\s+of\s+clusters\|resolution)\b |
| 9 | \bempiric(?:al\|ally)\b |
| 10 | \bconsistent\s+with\s+(?:previous\|prior)\s+(?:work\|studies)\b |
| 11 | \bfollowing\s+(?:previous\|prior)\s+(?:work\|studies\|protocol)\b |
| 12 | \bas\s+described\s+(?:previously\|elsewhere)\b |


evaluation\_foundbinary: any hit in DEFAULT\_PATTERNS["evaluation"]

Set to 1 if at least one regex in DEFAULT\_PATTERNS["evaluation"] matches the body text; otherwise 0.
This captures explicit cluster validation or model diagnostics (silhouette, DBI, CH, ARI/NMI, BIC/AIC, stability, modularity, etc.).
If the value is 0, then missing\_evaluation=1.

| # | Regex (verbatim) |
| --- | --- |
| 1 | \bsilhouette(?:\s+score\|\s+coefficient)?\b |
| 2 | \bdavies[-\s]?bouldin\b |
| 3 | \bcalinski[-\s]?harabasz\b |
| 4 | \bwithin[-\s]?cluster\s+sum\s+of\s+squares\b\|\bWCSS\b |
| 5 | \bbetween[-\s]?cluster\s+variance\b |
| 6 | \bcluster\s+stability\b\|\bstability\s+analysis\b |
| 7 | \bbootstrap(?:ping)?\b.\*\bcluster\b |
| 8 | \badjusted\s+rand\b\|\bARI\b |
| 9 | \bnormalized\s+mutual\s+information\b\|\bNMI\b |
| 10 | \badjusted\s+mutual\s+information\b\|\bAMI\b |
| 11 | \bfowlkes[-\s]?mallows\b\|\bFMI\b |
| 12 | \bjaccard\b |
| 13 | \bpurity\b |
| 14 | \bBIC\b\|\bAIC\b |
| 15 | \blog[-\s]?likelihood\b |
| 16 | \bmodularity\b |
| 17 | \bconductance\b |


tuning\_foundbinary: any hit in DEFAULT\_PATTERNS["tuning"]

Set to 1 if at least one regex in DEFAULT\_PATTERNS["tuning"] matches the body text; otherwise 0.
This captures explicit hyperparameter search language (grid/random search, Bayesian optimization, Optuna/Hyperopt, CV, etc.).
If the value is 0, then missing\_tuning=1.

| # | Regex (verbatim) |
| --- | --- |
| 1 | \bhyper[-\s]?parameter\s+(?:tuning\|optimization\|optimisation)\b |
| 2 | \bparameter\s+sweep\b |
| 3 | \bgrid\s+search\b |
| 4 | \brandom(?:ized)?\s+search\b |
| 5 | \bbayesian\s+optimization\b\|\bbayesian\s+optimisation\b |
| 6 | \boptuna\b\|\bhyperopt\b\|\bskopt\b |
| 7 | \bcross[-\s]?validation\b\|\bCV\b |
| 8 | \bnested\s+cross[-\s]?validation\b |
| 9 | \b(?:tuned\|optimized\|optimised\|selected)\s+using\b |
| 10 | \bwe\s+(?:tuned\|optimized\|optimised)\b |


missing\_paramsderived: params\_found empty

Computed as int(len(params\_found)==0).


missing\_justificationderived: justification\_found==0

Computed as int(justification\_found==0).


missing\_evaluationderived: evaluation\_found==0

Computed as int(evaluation\_found==0).


missing\_tuningderived: tuning\_found==0

Computed as int(tuning\_found==0).


missing\_reporting\_signalsderived: OR of missing flags

Computed as int(missing\_params or missing\_justification or missing\_evaluation or missing\_tuning).


parse\_errorsrow status: extraction failure

If XML parsing/extraction/analysis raises an exception, the row is still emitted with an error field set.
Downstream, year-level parse\_errors counts such rows.
